# Surprise response as a probe for compressed memory states

**DOI:** 10.1101/627133

**Authors:** Hadar Levi-Aharoni, Oren Shriki, Naftali Tishby

## Abstract

The limited capacity of recent memory inevitably leads to partial memory of past stimuli. There is also evidence that behavioral and neural responses to novel or rare stimuli are dependent on one’s memory of past stimuli. Thus, these responses may serve as a probe of different individuals’ remembering and forgetting characteristics. Here, we utilize two lossy compression models of stimulus sequences that inherently involve forgetting, which in addition to being a necessity under many conditions, also has theoretical and behavioral advantages. One model is based on a simple stimulus counter and the other on the Information Bottleneck (IB) framework. These models are applied to analyze a novelty-detection event-related potential commonly known as the P300. The trial-by-trial variations of the P300 response, recorded in an auditory oddball paradigm, were subjected to each model to extract two stimulus-compression parameters for each subject: memory length and representation accuracy. These parameters were then utilized to estimate the subjects’ recent memory capacity limit under the task conditions. The results, along with recently published findings on single neurons on the IB model, underscore how a lossy compression framework can be utilized to account for trial-by-trial variability of neural responses at different spatial scales and in different individuals, while at the same time providing estimates of individual memory characteristics at different levels of representation using a theoretically-based parsimonious model.

**Author summary:** Surprise responses reflect expectations based on preceding stimuli representations, and hence can be used to infer the characteristics of memory utilized for a task. We suggest a quantitative method for extracting an individual estimate of effective memory capacity dedicated for a task based on the correspondence between a theoretical surprise model and electrophysiological single-trial surprise responses. We demonstrate this method on EEG responses recorded while participants were performing a simple auditory task; we show the correspondence between the theoretical and physiological surprise, and calculate an estimate of the utilized memory. The generality of this framework allows it to be applied to different EEG features that reflect different modes and levels of the processing hierarchy, as well as other physiological measures of surprise responses. Future studies may use this framework to construct a handy diagnostic tool for a quantitative, individualized characterization of memory-related disorders.

## Introduction

There is abundant experimental evidence from behavioral studies on the finite capacity of recent memory [1, 2]. Limitations on memory size, the rate of writing and reading from memory, and the amount of information that can be transferred in a unit of time (i.e., channel capacity [3]) can all impose boundaries on the history length and accuracy of memory representations. Depending on task requirements, these constraints may lead to compressed representations of the observed past. Compressed representations are also associated behaviorally [4–7] and theoretically [8–10] with more efficient learning and generalization. Compressed representations currently cannot be measured directly in the human brain, but nevertheless can still impact the electrophysiological responses to each stimulus in a sequence as a function of the representation of preceding stimuli (for a similar methodology see [11]).

Surprise (alternatively called *surprisal* [12, 13] in the predictive coding literature) is one possible way to study compressed memory states. Intuitively, the lower the accuracy of past stimulus representations, the more future variations in the stimuli will go unnoticed, causing lower surprise responses. Conversely, the more accurately past stimuli are represented, the more surprising small variations will be in future stimuli. This intuition is consistent with behavioral and neural data [14–16] showing larger error (surprise) responses to deviants in music in trained musicians as compared to non-musicians.

The P300 (or more specifically, the P3b [17]) is a well-known event-related potential (ERP) component in electroencephalography (EEG) measurements [18] evoked by rare, task-relevant events, and is typically observed in oddball paradigm experiments in which the oddball is the target stimulus. The average P300 response to an oddball stimulus is known to have a higher amplitude the lower the probability of the oddball [19]. In addition, more complex models have indicated a relationship between the P300 amplitude and certain definitions of trial-by-trial surprise (see supplementary S1 Text). Furthermore, working memory studies [20–22] have shown that working memory load causes a decrease in the amplitude of the P300, which is consistent with the notion that lesser memory capacity leads to a lower surprise response. For all these reasons the P300 seems a good candidate to serve as a physiological surprise probe for compressed memory states.

Under an auditory oddball paradigm, we tested two models of compression-dependent surprise, both involving lossy compression. This lossy compression should be contrasted with a compression that enables remembering each stimulus in a sequence, i.e., with lossless compression that allows perfect reconstruction of the past stimuli. While this kind of precise memory could be necessary in behavioral memory tests such as recall tests, it is less relevant for prediction in many cases [23, 24]. The hypothesis was that the surprise responses would correspond to a representation in which the forgotten details are those that are less relevant for predicting the next stimulus in the sequence. The first model we consider is a naive oddball count (NOC) model. It estimates the surprise associated with a given stimulus in a sequence based on the number of oddball stimuli in a given time window into the past. It is lossy in the sense that the order of the elements is not preserved in the representation, but is also optimal in the sense that it preserves the exact statistics of the past oddball sequence required for predicting the next stimulus.

Although optimal prediction in the oddball sequence requires that the exact number of preceding oddball tones be remembered, it is more plausible that this number is only roughly estimated and remembered. This hypothesis is formalized mathematically in the second model [25] that draws on the Information Bottleneck (IB) framework [26] (see Fig. 1 and Methods). Importantly, the IB shows that even with an approximation of the oddball count, good prediction accuracies of the next stimulus can be achieved [25].

**Fig 1.**
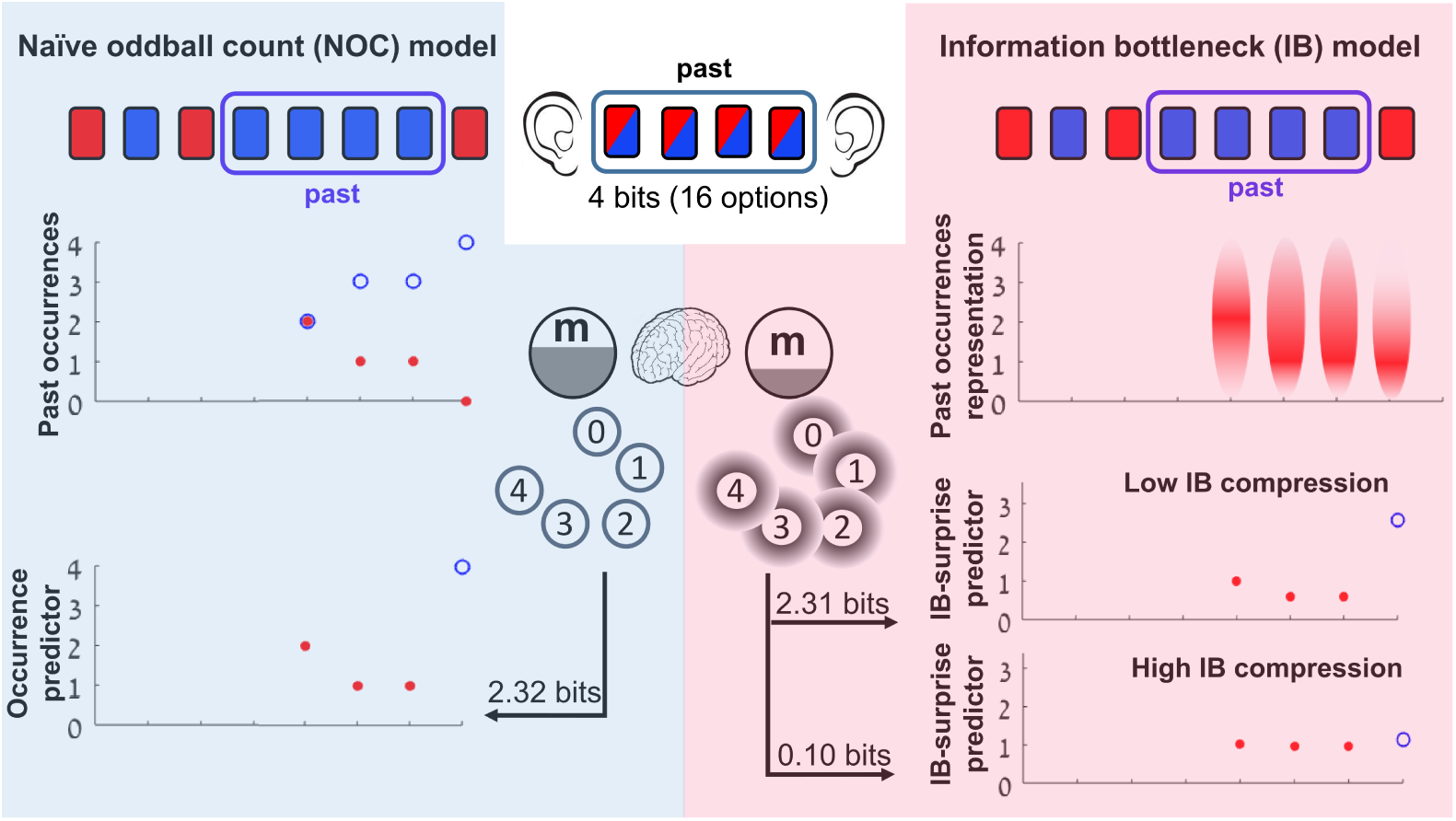
The naive oddball count (NOC) and the IB lossy compression models of the oddball sequence. An illustration of the two compression models for the case of a past length of size *N* = 4 is shown. To be able to code any of the 16 possible sequences of 4 tones in memory that the subject heard, 4 bits of memory would be needed. (**Left**) The NOC model only keeps the number of oddball occurrences in the previous window in memory; i.e., the minimal sufficient statistic (see Methods). The upper plot shows the number of oddball tone occurrences *n* (filled red circles) and the number of standard tone occurrences *N*−*n* (empty blue circles) in the previous window for each trial, starting from trial *N* + 1. All plots are aligned to the stimulus sequence. The occurrence predictor (bottom plot) on each trial is *n* if a standard tone was played and *N*−*n* if an oddball tone was played. To be able to code the 5 alternatives of the past in memory, 2.32 bits of memory are required. (**Right**) The IB model keeps a fuzzy representation of the oddball counter in memory, which requires less memory usage than the NOC model. The upper plot illustrates a fuzzy representation *m* of the oddball occurrences *n* in the previous window, as given by IB for a high compression case. The darker red represents higher *p*(*m*|*n*) probability. The two lower plots show two IB predictors of different compression levels (0.1 bits for the bottom plot, 2.31 bits for the middle plot). The surprise level on each trial is defined as -Σ*_m_ p*(*m*|*n*) log *p*(*next tone*|*m*) where the probabilities are defined by the IB solution for a specific compression level.

In a trial-by-trial analysis, we show that both the NOC and the IB models achieve significant goodness-of-fit to the P300 response, providing support for the lossy compression hypothesis. While the models’ performance was comparable, there was a notable difference in the estimated memory usage (here termed effective capacity) which was significantly lower for the IB model. Hence, while both lossy compression models are consistent with the EEG results, the IB model emerged as a more parsimonious model in terms of memory utilized for the task. Overall, these results demonstrate how lossy compression can provide a framework for analyzing trial-by-trial responses (see Rubin et al. for single cells [25]), and show how these responses can be used to obtain an estimation of recent memory usage.

## Results

We presented 17 subjects (age 31±10; 10 females) with a two-tone auditory oddball sequence (see Fig. 2a), containing blocks with high-tone probabilities of 0.1 to 0.5. To encourage the subjects to attend to the tones and anticipate the next tone, they were asked to press a button as quickly as possible each time they heard the high tone. Throughout the experiment their EEG response was recorded (see Methods).

**Fig 2.**
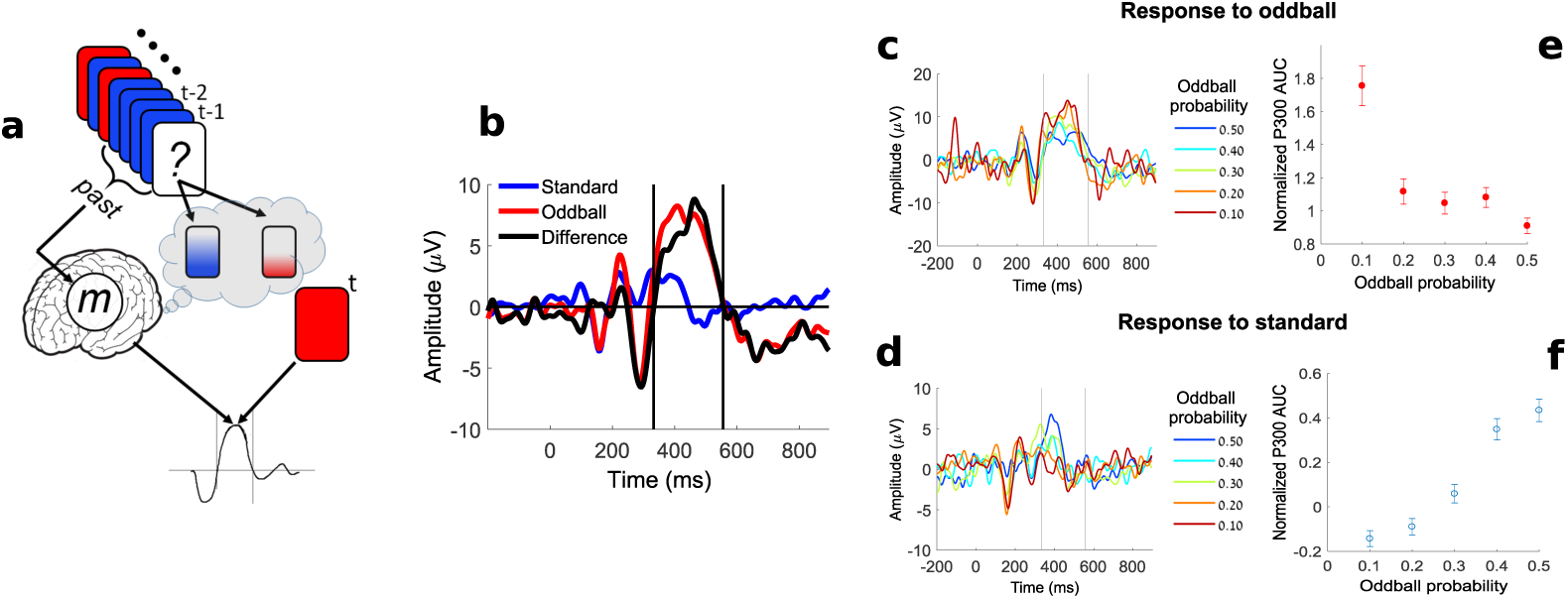
The model for the oddball experiment and the EEG surprise measure. (**a**) A two-tone oddball sequence is presented to the subject. The previous stimuli are processed and the subject holds an internal representation *m* of the past in memory. Based on *m* the subject holds a prediction of the next tone (high vs. low). The subject’s response to the tone at time *t* is dependent on the tone type and the incomplete memory *m* of the past. A fixed window size *N* of the past is considered on each trial (in this illustration *N* = 4). (**b**) The definition of the P300 area-under-the-curve (AUC): event-related potentials averaged over all oddball (red) and standard (blue) trials in electrode Cz are shown for a representative subject. The difference between the oddball and standard curves (solid black) was used to determine the zero-crossing points around the P300 peak (dashed vertical lines). The P300 AUC per trial was defined as the area between these two time points on each trial. (**c**), The average trace of all standard trials in each block is shown color-coded (for the same subject as in (b)). The oddball probability (OP) of each block is shown in the legend. (**d**), The average trace of all oddball trials in each block is shown color-coded (as in (c)). (**e**), The mean normalized AUC of all oddball trials in each block averaged over all subjects are plotted as a function of the corresponding oddball probability of that block. The error bars indicate the standard error of the mean (SEM). (**f)** The same as (e) for standard trials.

### The P300 area-under-the-curve as a measure of physiological surprise

The P300 amplitude is known to be inversely dependent on the oddball probability (OP) [27], a necessary condition for it to be used as an EEG surprise feature (as done in several studies; see supplementary S1 Text). The area-under-the-curve (AUC) is a sum of amplitudes; therefore, it is less prone to noise than the P300 amplitude and may serve as a better feature for trial-by-trial analysis.

We used an automatic procedure to define the P300 AUC time range for each subject by first finding the maximal peak in the range of 300 to 500 ms (after the stimulus onset) in the mean oddball-to-standard difference curve at electrodes Cz and Pz (Fig. 2b). Since electrode Cz showed higher maximal peaks on average across subjects (5.37 *µ*V for Cz vs. 4.48 *µ*V for Pz), the remainder of the analysis was performed on the Cz channel. The time points between which the AUC was calculated were defined as the zero-crossing points of the oddball-to-standard difference trace (Fig.2b). The P300 AUC for each trial was calculated using these time points.

In Fig. 2c and 2d we show the average response signal for oddball and standard trials for each of the oddball probabilities for a representative subject. As reported previously [27], when the OP is decreased, the magnitude of the response increases for the oddball trials (Fig. 2c). In Fig. 2d we show that there was also a decrease in the magnitude of the P300 in response to standard trials with decreasing OP.

For the multi-subject analysis below, we normalized the AUC values of each subject by dividing them with the AUC of the mean oddball-to-standard difference trace of that subject (black curve in Fig. 2b). In Fig. 2e we show the mean normalized AUC averaged over all subjects as a function of the oddball probability. The AUC for oddball and standard trials exhibited a similar trend as the known dependency of the P300 amplitude on the oddball probability.

### Single-subject analysis of single-trial P300 data using the NOC and IB models

Fig. 3a depicts the dependency of the single-trial P300 AUC on the number of oddball occurrences *n* (as defined by the NOC model, Fig. 1) in the preceding *N* = 11 trials in the sequence, for a representative subject. The optimal *N* was determined by the *N* value with the maximal goodness-of-fit, calculated using weighted linear regression (see Methods). High tone trials are marked by solid red circles and low tone trials are indicated by empty blue circles.

**Fig 3.**
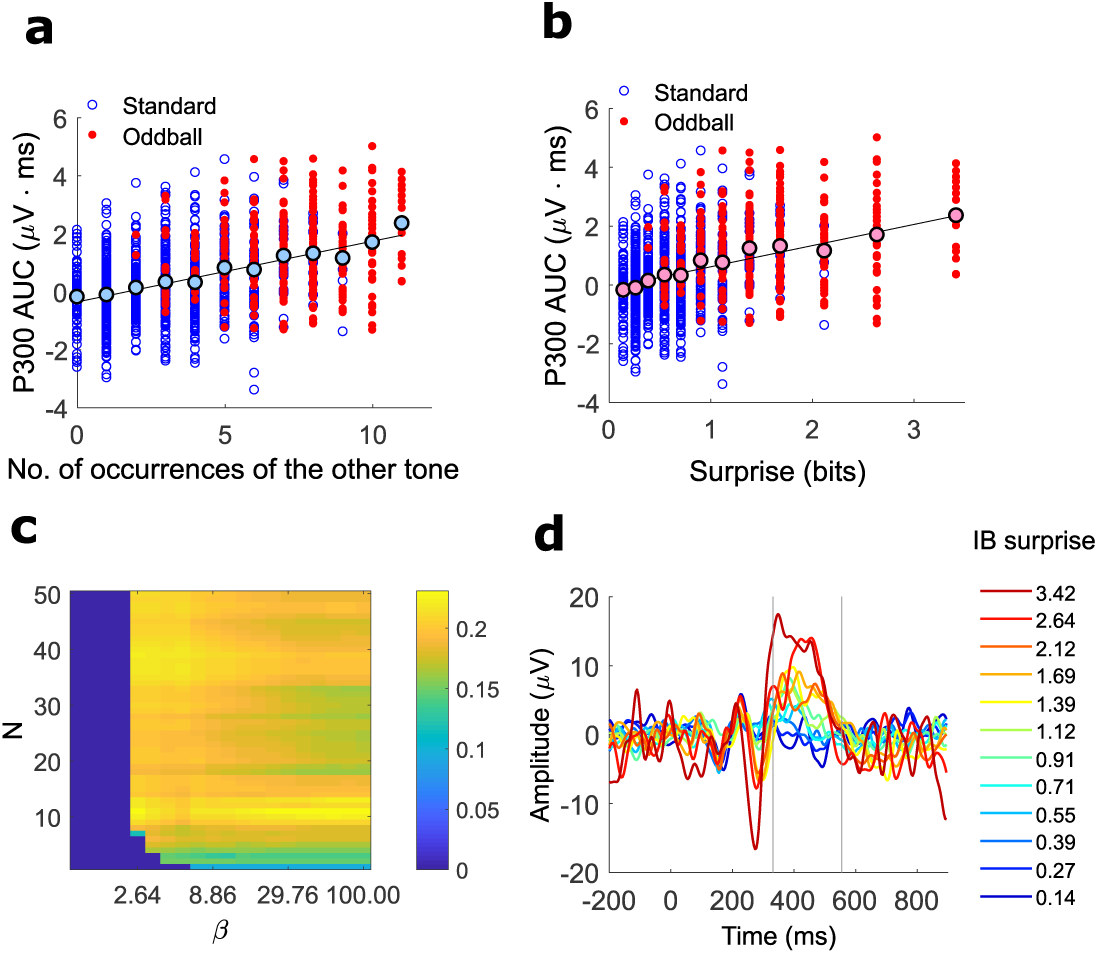
Subject-specific compression parameters extracted by single-trial analysis. (**a**) Single-trial P300 AUC responses of a representative subject to standard tones (blue empty circles) and to oddball tones (red, filled circles) as a function of the number of occurrences (*n*) of the opposite tone in the preceding sub-sequence of *N* = 11 tones (the fitted *N* for that subject). The black-edged filled circles show the average response for each *n*. Single trials: weighted-*R*^2^=0.230, one-sided permutation test for the *R*^2^, 1000 permutations, p-value*<*0.001, 1145 data points. Mean values: *R*^2^ = 0.934 p-value=3.14×10^−7^, 12 data points. (**b**) Single-trial P300 AUC responses to standard tones (blue, empty circles) and to oddball tones (solid red circles) as a function of the optimal IB surprise predictor (*N* = 11, *β* = 48.33) for the same subject as in (a). The black-edged solid circles show the average response for each surprise value. Single trials: weighted-*R*^2^=0.231, one-sided permutation test for the *R*^2^, 1000 permutations, P-value¡0.001, 1145 data points. Mean values: *R*^2^ = 0.939 p-value=2.22×10^−7^, 12 data points. (**c**) color-coded weighted-*R*^2^ values of the linear regression analysis for all tested IB compression parameters, for the same subject as in (a,b). The vertical axis denotes the memory window length *N*. The horizontal axis denotes the representation accuracy parameter *β*. The pair of model parameters (*N, β*) that achieved the highest *R*^2^ value was used for the remainder of the analysis. (**d**) Surprise-related analysis (SRA). The single-trial waveforms were averaged by the IB surprise associated with each trial and are shown color-coded according to the surprise level (shown in the legend). The average SRA waveforms exhibited a gradual increase in the P300 magnitude with the IB surprise level.

The average over all the trials with the same number of occurrences *n* (light blue, black edged circles) showed an increase with *n*; i.e., the higher the number of occurrences of the opposite tone in the previous window, the larger the P300 response to the current tone. Note that this plot combines two experimental phenomena: response increase with the rarity of a stimulus (novelty detection) and response decrease the more a stimulus was repeated (adaptation), both of which were characterized by the variable *n*.

Fig. 3b depicts the dependency of the single-trial P300 AUC on the IB surprise generated by the IB model with *N* = 11 and *β* = 48.33, for the same subject as in Fig. 3a. The optimal *N* and *β* are determined by the pair of (*N, β*) values with the maximal goodness-of-fit, calculated using weighted linear regression (see the weighted-*R*^2^ map in Fig. 3c and Methods SI). The mean AUC values per surprise showed a gradual increase with the theoretical surprise value; i.e., the higher the surprise, the larger the P300 response. For examples of more subjects see supplementary S3 Fig For a qualitative and quantitative comparison to other surprise models, see supp. S1 Text and supp. S5 Fig

Another way to assess the dependency of the P300 on surprise is by averaging the traces of all the trials with the same theoretical surprise value, here termed surprise-related analysis (SRA) (Fig. 3d). The average trace of each surprise value is plotted in a different color, where the color code bar indicates the surprise value. As reflected in the mean AUC values, the P300 response gradually increases as the theoretical surprise increases (for an analysis of the oddball trials alone see supplementary S5 Fig).

Note that the SRA is essentially a generalization of the more standard ERP analysis for oddball sequences (e.g. as in Fig. 2b), in which all oddball events are considered as surprising and are averaged together, and all standard events are considered non-surprising and are averaged together, thus masking the gradual increase in the amplitude. The SRA is also a generalization of the plots presented in Fig. 2c,d.

### A multi-subject comparison of the NOC and the IB models

For all subjects, the maximal weighted-*R*^2^ obtained with the IB model was similar or slightly higher than the maximal weighted-*R*^2^ obtained with the NOC model (see a subject-by-subject comparison in supplementary S2 Fig). To examine the combined data of all subjects, we used the normalization of the AUC on the y-axis, as defined for Fig. 2e,f. For the NOC model we had to use a normalization of the occurrences for the x-axis since every subject had a different value for the memory length *N*. Therefore we normalized the number of occurrences *n* with *N* (for each subject) to define a running probability (RP). The IB surprise, on the other hand, provides a universal scale for all subjects and needs no normalization. The mean responses of all subjects are plotted in Fig. 4a,b. A linear trend was observed in both (NOC: *R*^2^=0.498, IB: *R*^2^=0.471). The plots in the insets show the corresponding averaged responses. While the fitting accuracies were comparable, it should be noted that the IB surprise accounted for a wider range of P300 responses (as can be seen from the range of the vertical axes in the insets of Fig. 4a,b).

**Fig 4.**
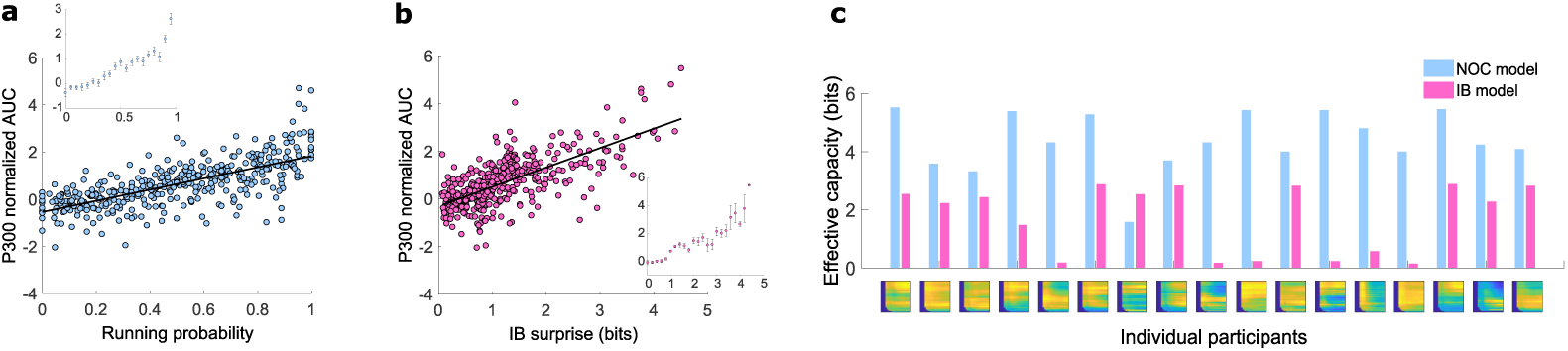
Multi-subject comparison of the NOC and IB models. (**a**) The mean normalized AUC of each subject is plotted as a function of the running probability (RP). The RP on each trial is defined as *p* = *n/N,* where *n* is the number of occurrences of the opposite tone (with respect to the current tone) in the preceding *N* tones in the sequence, and *N* is the fitted model parameter. *R*^2^ = 0.498, 409 data points, error DOF= 407, F-statistic vs. constant model: 404, p-value=7.28×10^−63^ (**b**) The mean normalized AUC of each subject is plotted as a function of the IB surprise. *R*^2^ = 0.471, 445 data points, error DOF = 443, F-statistic vs. constant model: 394, p-value = 0. (**c**) The mean responses across all subjects as a function of the RP, calculated using the data presented in (a). The error bars indicate the SEM. (**d**) The mean responses across all subjects as a function of the IB surprise, calculated using the data presented in (b). The error bars indicate the SEM. (**e**) Each pair of bars shows the estimated effective capacity (in bits) for the optimal NOC model (blue bars) and for the optimal IB model (red bars) for each subject. The effective capacity for the NOC model is defined as the number of bits required to represent all possible values of occurrences from 0 to *N*; i.e., *log*_2_(*N* + 1). For the IB model the estimated capacity is defined as *I*(*X*; *M)*; i.e., the mutual information between the past variable *X* = [0..*N*] and the representation variable *M* defined by IB. The colored map under each pair of columns shows the IB *R*^2^ map of that subject (as in Fig. 3c).

In order to quantify the amount of compression under each model, we calculated the amount of memory (in bits) required for the past internal representation for each subject, given the fitted model parameters (*N* for the NOC model and *N, β* for the IB model). This is effectively an estimation of the individual’s memory capacity that was relevant to the task, and thus was dubbed *effective capacity*. Fig. 4c shows a subject-by-subject comparison of the effective capacity. Although the weighted-*R*^2^ was fairly similar for the two models (see also supplementary S2 Fig), in terms of the amount of utilized memory, the IB model suggests an account for the data by a much more memory-efficient representation.

## Discussion

The estimation of memory capacity has been addressed by numerous studies that have typically used behavioral measures [1, 3, 28–30]. Here we suggest a novel, indirect approach for estimating recent memory capacity by utilizing the dependency of neural responses to sequential stimuli on the internal representation of past stimuli. While this framework is very general, we successfully demonstrated the concept using a well-studied experimental paradigm in the ERP literature known as the oddball paradigm and the P300 ERP component. By using a compression-dependent surprise, a highly parsimonious model originally developed and used for explaining single-neuron responses [25], we were able to account for single-trial variations in the P300 response. This model also provided a quantitative estimate of the effective recent memory capacity for each subject, which may be thought of as the amount of utilized memory that was relevant to the task. By defining a task that is not explicitly memory-dependent, we unconstrained this amount of memory and allowed natural forgetting of the stimulus sequence. The framework included three types of forgetting: (1) forgetting the order of the sequence (2) forgetting the distant past beyond the previous *N* elements and (3) diminished accuracy of the past stimuli representation. We posited that subjects forget as many of the stimulus details as possible, as long as the task performance is not impaired above some individually set threshold. This hypothesis, and the optimal compression defined by the three types of forgetting above, gained evidence from the trial-by-trial P300 correspondence with the NOC model, and to an even greater extent with the IB model. Importantly, the generality of the framework allows it to be applied to various surprise (or prediction error) signals, thus yielding a memory capacity estimation at different modes and levels in the hierarchy of stimulus processing (see supplementary S4 Fig for the correspondence between surprise and response time).

To situate our results in the context of the current memory literature, the fuzzy representation of the number of oddball occurrences (as illustrated in Fig. 1 right) has much in common with the interference approach to working memory (WM) [31]. According to the IB framework these “interfering” representations are the optimal form of forgetting in limited capacity conditions in a given task. Since WM is also considered task dependent [32], it implies an association between the memory capacity discussed here and WM. It is also worth noting that in both of the compression models presented here, less probable stimuli elicit higher surprise responses, which is consistent with the “novelty-gated encoding” assumption of interference models [31]. However, WM capacity is typically considered to be an inherent trait of each individual, whereas the effective capacity we estimated here should be seen as memory devoted to the specific task under the current state of the subject (that can be affected by motivation, attention, fatigue etc.). Another important difference is that typically WM is thought to contain under 10 elements which are represented rather accurately in memory [33]. In contrast, here we consider the effect of up to 50 elements in the past. These elements are not remembered accurately according to the optimal (lossy) compression models; rather, the relevant information in them affects the response to the current stimulus. Thus the plausible link between WM and the relevant recent memory discussed here should be tested in future experiments.

Another natural and intriguing link is to the predictive coding framework [12, 34]. Within this framework it is hypothesized that the cortex constantly generates predictions of incoming stimuli, and responds to deviants from these predictions by a prediction error signal. More specifically, in hierarchical predictive coding theories [12, 35], the P300 has been linked to deviants from higher-order expectations [36–38]. We expand on these results by reporting a more subtle P300 response to both deviants and standards (Fig. 2e-f, Fig. 3d), and also suggest a computational framework to experimentally examine and characterize prediction error signals at various levels of the hierarchy, by identifying the past length and resolution parameters (i.e., *N, β*) that best fit the data. There is also a compelling relation between the resolution quantified by *β* and the precision expectation model of attention [**?**] which should be explored in future studies.

We presented the results of a phenomenological model of an optimal strategy under memory limitations. The advantage of this type of theoretical modeling is that it allows for quantitative predictions with very few assumptions and parameters. However, the specific mechanism by which the sequence of stimuli is transformed into a concise memory state, the way in which this memory state affects the neural response to a new stimulus or how a new memory state is created are beyond the scope of this paper. Rather, the type of modeling presented here poses constraints on any such possible mechanisms. Nevertheless, the P300, a well-studied ERP component, and the very common oddball paradigm used in both human and non-human studies, makes it possible to relate the results to known underlying mechanisms. Importantly, Rubin et al. [25] showed that the IB model was able to explain the firing rate variability of early single-neuron responses in the primary auditory cortex to oddball sequences. This provides striking evidence for a lossy type of compression affecting responses as early as at the single neuron level. The relationship between these single-neuron results and our results on the higher-scale P300 ERP component calls for further investigation; however, the same surprise model applied to both phenomena provides an intriguing link that may shed more light on these multi-scale phenomena of novelty on one hand, and attenuated responses to repeated stimuli on the other. With regard to the function of these phenomena, both the P300 and the single neuron prediction-error responses have been associated with memory update processes [39, 40]. The combination of compression-dependent prediction-error responses with memory-update mechanisms that depend on these deviant responses may hint at a possible mechanism for the formation of compressed representations [41, 42].

Although the underlying mechanisms currently remain elusive, the literature on the P300 still allows us to discuss several possible implications deriving from this model. The P300 is known to be related to numerous factors such as attention [43], mental workload [44], age [45] [46] and even neurological pathologies [47]. The NOC and IB compression models provide a method to identify individual memory characteristics in a handy way that is simple enough to be applied even on children [48] or individuals incapable of complex tasks. This may contribute to developing quantitative diagnosis of abnormalities related to short-term processing in various pathological conditions.

## Methods

### Subjects

Twenty healthy participants (age 31±10; 12 females) participated in the experiment, of whom three were excluded according to criteria described below. All participants were recruited via ads at Ben-Gurion University. The experimental procedures were approved by the institutional ethics committee at Ben-Gurion University. Prior to the experiment, the participants signed a written consent form after being properly informed of the nature of the experiment. All participants were compensated for their participation.

### Experimental protocol

Before the experiment, the participants were presented with a short example of a two-tone sequence of 14 tones, out of which 2 were high tones. They were instructed to press the space bar as quickly as possible each time they heard the high tone. The aim of the example was to let the subjects know which tone was the high tone and which was the low, and to make sure they understood the task. If they did not press the space bar each time the high tone was played, the example was played again. If they still failed they were excluded from the database (one subject).

The experiment was made up of 5 blocks of a two-tone sequence, each containing 240 trials (tones). Each block had a different oddball probability [0.1, 0.2, 0.3, 0.4, 0.5] and the tone for each trial was drawn independently from a Bernoulli sequence. On each trial, a low tone (with a frequency of 1000 Hz) or a high tone (1263 Hz) was played for 125 ms. The participants had to detect whether it was the high or the low tone and were required to press the space bar as quickly as possible if it was the high tone. They did not receive feedback on their performance and the next tone was played with a fixed inter-stimulus interval (ISI) of 1 sec whether they were right or wrong. Their reaction time (RT) for each button-press was measured (with the exception of one subject for whom RT data was not measured for technical reasons). For each subject a new sequence was drawn and the order of the blocks was randomized. Between blocks the participants were given a break and could continue to the next block whenever they wished. Subjects were not informed about how the sequences were generated. The experiment was realized using Matlab R2015a with the Psychophysics Toolbox extension [49–51].

### Data acquisition

EEG was recorded (bandpass filter: 0.1–60 Hz 8th order Butterworth, notch filter: 48–52 Hz 4th order Butterworth, 256 Hz sampling rate) using a g.HIamp amplifier and a g.GAMMAsys cap (g.tec, Austria). For 8 subjects the EEG was recorded from the following electrode positions: Fz, Cz, Pz, P3, P4, PO7, PO8, Oz, and left and right ear electrodes. For the remaining 11 subjects the EEG was recorded using 61 electrodes positioned according to the extended international 10–20 system, and left and right ear electrodes. The anterior midline frontal electrode (AFz) served as the ground. The active electrode system g.GAMMAsys transmits the EEG signals at an impedance level of about 1 kΩ to the amplifier.

### EEG data preprocessing

EEG data were analyzed using Matlab R2017a (Mathworks) with the EEGLAB [52] toolbox (v14.1.1b) and the ERPLAB plugin (v6.1.4). Each participant’s EEG data were bandpass filtered (0.1–30 Hz), and re-referenced to the average of the two ear electrodes. Subsequently, epochs of −200 ms to 900 ms around the presentation of the tones were extracted from each trial. We should first note that data preprocessing and analysis procedure was conducted with the goal in mind to have an automatic, fast and reliable procedure, for the sake of a possible later usage within a diagnostic or treatment tool (e.g. in a neurofeedback treatment). To this end, we used a standard automatic artifact-cleaning procedure based on the artifact subspace reconstruction method [53, 54], by applying the clean rawdata extension for EEGLAB with the parameters arg flatline: 5, arg highpass: [0.25 0.75], arg channel: 0.7, arg noisy: 4, arg burst: 5, arg window: ‘off’. We then used independent component analysis (ICA) ([55] and components classification for removing artifactual components using the MARA plugin for EEGLAB [56, 57]. In MARA, we modified the threshold of artifact probability p artifact to 0.05 (instead of the default 0.5), so that components with a p artifact higher than 0.05 were removed. On average, 39% of the components were removed for the 64-electrode datasets (in the 8-electrode datasets there was only one component overall that was removed).

We first calculated the mean oddball and standard responses for each subject at electrodes Cz and Pz. Since electrode Cz showed higher P300 amplitudes on average across subjects (5.37 *µ*V for Cz vs. 4.48 *µ*V for Pz in the difference between oddball to standard), the rest of the analysis was performed on the Cz channel. Having a clear P300 peak (by visual “eyeballing”) in the averaged ERP was determined as an inclusion criteria for the study, therefore subjects for whom there was no clear peak in the mean oddball response compared to the mean standard response in the relevant time range (300-500 ms) were excluded from the rest of the analysis (two subjects, see supplementary S1 Fig for the single-subject ERP plots of all subjects). In order to define the time points for the calculation of the single-trial P300 area-under-the-curve (AUC), we calculated the difference between the mean oddball response and the mean standard response (Fig. 2b). We then automatically extracted the time point *t*_*peak*_ of the maximal amplitude between 300 ms to 500 ms in this difference trace. Around *t*_*peak*_, we extracted the two time points *t*_1_, *t*_2_ where the trace crossed the horizontal axis. The single-trial P300 feature was defined as the area-under-the-curve (AUC) between *t*_1_ and *t*_2_ for each trial. It is worth noting that we used the AUC rather than the peak amplitude because it demonstrates a similar dependency on the oddball probability as the P300 amplitude (see Fig. 2e), but is theoretically expected to be a more robust feature of the single-trial P300 than the amplitude since it is composed of a sum of amplitudes. The custom Matlab routines developed for the study are available from H.L.A.

### The naive oddball count (NOC) model

The mathematical details are given in the sections below. In short, the NOC model is based on the minimal sufficient statistics (MSS) of the oddball sequence (Fig. 1 left) in the following way: the predictor of the NOC model is constructed by first calculating the number of oddball tones and the number of standard tones in a fixed window of size *N* of the preceding sequence (Fig. 1 left, upper plot). *N* is the only parameter of the model and represents the memory length affecting the current stimulus response; i.e., how far into the past the stimuli are counted. The response for the tone on each trial is then modeled by the number of occurrences of the opposite tone in the past window; e.g., if the high tone was played on a trial, the number of occurrences of the low tone in the past window is taken as the predictor value for that trial (Fig. 1 left, bottom plot). It should be noted that this simple model suggests a unified theoretical explanation for two well-known multi-scale phenomena: the decreased response to repeated stimuli and the increased response to novel or rare stimuli. In the following sections we give the mathematical arguments for the optimality of this predictor.

### Maximum likelihood estimation and sufficient statistics in the oddball paradigm

In the field of parameter estimation, a sufficient statistic is a function of the data that summarizes all the relevant information in the data for estimating a specific parameter (for a formal definition see [58] p.35). For example, the empirical mean of a sequence of random numbers is a sufficient statistic for estimating the mean of the distribution.

Here we consider a typical oddball sequence *z*_1_, *z*_2_, *…z*_*N*_, *z*_*i*_ ∈{0, 1} generated using a Bernoulli probability distribution:

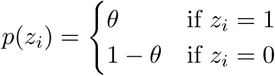

where *θ* is the oddball probability, also called the Bernoulli parameter. The *z*_*i*_’s are referred to below as samples. By definition, a sufficient statistic for the Bernoulli parameter *θ* is the empirical mean of the sequence: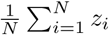. Moreover, it can be shown that the empirical mean is a *minimal* sufficient statistic (MSS), in the sense that given this number, no additional information about the sequence (e.g. the order of the elements) can improve the estimation of the oddball probability. For a formal definition of a minimal sufficient statistics see [58] p.37. It is worth noting that for the Bernoulli sequence, the maximum likelihood (ML) estimator of *θ* is tightly related to the MSS, since 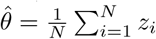, where 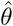 denotes an estimator of *θ*.

### Prediction in the oddball sequence

We now consider a “past” sequence *z*_1_, *z*_2_, *…z*_*N*_ and a prediction of the “next” element *z*_*N*+1_ (denoted by *y*). Given the ML estimator 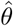 defined in the previous section, an optimal prediction for y is given by the Bernoulli distribution with 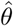 as the Bernoulli parameter, i.e.:

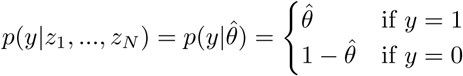

Note that 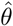 is simply the number of oddball occurrences in the *N* previous elements, divided by *N*. The meaning of this well-known result is that simply keeping track of the number of oddball occurrences in the past sequence is a powerful predictor of the next tone.

### The model predictor

As described above, the predictor of the NOC model is constructed by first calculating the number of oddball tones and the number of standard tones in a fixed window of size *N* of the preceding sequence. The response for the tone at each trial is then modeled by the number of occurrences of the opposite tone in the past window (see Fig. 1 left).

Formally the predictor is defined as:

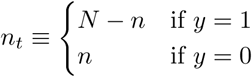

### The information bottleneck (IB) model

The concept of minimal sufficient statistics can be generalized to cases with limited memory resources. As mentioned above, given a minimal sufficient statistic for a parameter of a probability distribution, no additional information on the samples can improve the estimation of that parameter. On the other hand, it contains the minimal information for an optimal estimation; i.e., any further decrease in that information will result in a degraded estimation.

In cases of limited memory capacity, some information loss may be a necessity. Given a certain information loss, what is the accuracy that can be achieved when predicting the next element? The answer is provided by the IB method [26]; At each level of compression (information loss) it provides an upper bound on the accuracy that can be achieved in the prediction of a target variable (in our case the next tone in the sequence).

the IB model [25] uses the IB method to define a compression-dependent predictive surprise. Predictive surprise is defined as: [59–62]

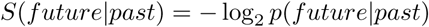

In the case of the oddball sequence the next element is considered as the *future*, and the *past* is the sequence of preceding elements to be considered. The probability *p*(*future*| *past*) is usually called the *subjective probability* since it is an estimation of the true probability from which the sequence was generated. While there is a solid rationale for the use of the log function [62, 63], different models of predictive surprise usually differ in the method employed for subjective probability estimation (see supplementary S1 Text). However, typically these models implicitly assume that the past is represented accurately in the brain.

Rubin et al. [25] considered the consequences of *subjective past*, to take into account the common case of limited memory capacity. As described above, an optimal representation of the past for the goal of predicting the next element is the sufficient statistic 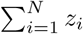 In cases of limited memory the IB method shows that the optimal compression is a representation with degraded precision (e.g., the subject only remembers there were “about” 4 oddball tones in the previous 10 tones). For these cases, the subjective surprise is not determined by the true past, but rather by its internal compressed representation *m*: *S*_*m*_(*future*)≡ −log *p*(*future*|*m*) and the probability of having a specific representation *m* is given by *p*(*m*|*past*). Both the probabilities *p*(*m*|*past*) and *p*(*future*|*m*) are determined by the optimal compression provided by the IB method. The mathematical details of the model are given in the following sections, as well as in [25].

### The IB method

Given a random variable *X* representing the samples or a function of the samples, a target variable *Y,* and a joint distribution *p*(*x, y*), *x* ∈*X y* ∈ *Y,* the IB method defines a compressed representation of *X* denoted by *M* that is most informative about the target variable *Y*. In our case *y* is the next element in the oddball sequence, *x* is the number of oddball occurrences in the previous *N* elements (i.e. 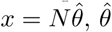 similarly as defined above), and *m* is a compressed representation of that number. Note that we defined the past *x* as 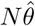 (rather than any other function of the past sequence) since we already know that 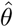 is an optimal compression of the sequence, and we are interested now in a further compression. Also note, that although *x* represents the “past” and *y* represents the “future”, the joint probability *p*(*x, y*) is assumed for simplicity to be stationary. This assumption can be relaxed by empirically estimating *p*(*x, y*) and updating it along the sequence.

The amount of compression of *X* is quantified by the mutual information between *X* and *M,* denoted by *I*(*X*; *M)*, whereas the information that *M* carries on the target variable *Y* is quantified by *I*(*M*; *Y)*. Ideally one would want to compress *X* as much as possible (to use minimal memory resources) while keeping the maximal amount of information on *Y* for optimal prediction accuracy; i.e. minimizing *I*(*X*; *M)* while maximizing *I*(*M*; *Y)*. Since there is a trade-off between the two, the optimal solution is determined by the minimum of the Lagrangian:

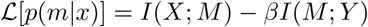

where *β* is the trade-off parameter, controlling the balance between the compactness of the representation and the preservation of relevant information; i.e., the lower the *β* the stronger the compression. For each *β* we obtain an optimal solution for *p*_*β*_ (*m*|*x*) and *p*_*β*_ (*y*|*m*) which satisfy:

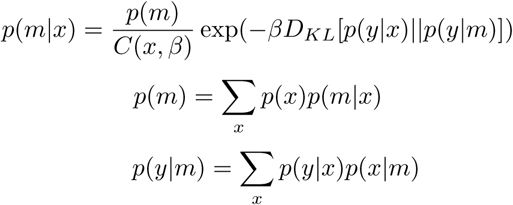

where *C*(*x, β*) is a normalization function, and *D*_*KL*_ is the Kullback-Leibler divergence [58] which is a distance measure for probability distributions. This set of self-consistent equations also determines an iterative algorithm to find the optimal solution for any given *p*(*x, y*) and *β*. For further details on the IB principle and algorithm see Tishby et al. [26].

### The model predictor

We denote by *y*_*t*+1_ the next element, and by *x*_*t*_ the minimal sufficient statistic of the *N* preceding elements: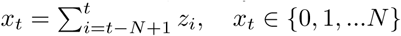, *x*_*t*_ ∈ *{*0, 1, *…N}*. Thus the IB predictive surprise is defined as:

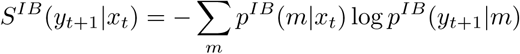

where *m* denotes a compressed representation of *x*_*t*_ and we summarize over all possible representations with their respective probability.

Note that we do not need to explicitly determine a specific compressed representation (although it can be done, see agglomerative IB [64]). As specified in the previous section, to use IB we only need the joint probability distribution of the past *x*_*t*_ and the future *y*_*t*+1_. To calculate this probability we assume that the subject has no prior knowledge of the sequence; therefore we use a uniform prior on the oddball probability which yields the following joint probability distribution:

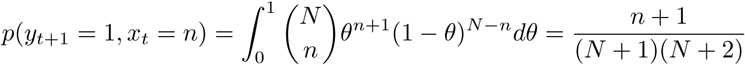

The IB surprise *S*^*IB*^(*y*_*t*+1_|*x*_*t*_) defined above is the subjective surprise level on each trial. It is dependent on two parameters: the memory length *N* and the compression parameter *β*. Note that in Fig. 1 *y*_*t*+1_ is simply referred to as the *next tone*, and for brevity, *n* denotes the random variable of the past (denoted here by *x*_*t*_), as well as the actual values obtained by *x*_*t*_.

By comparison with the NOC model, whereas in the NOC predictor the number of past oddball occurrences is accurately kept in memory (e.g. 2 oddball tones in the previous 4 tones), in the IB predictor this number can be represented in memory inaccurately, (e.g. 2 oddball tones in the past may be represented by a high probability for *n* = 2, but there is also some probability for other occurrences (0,1,3,4), as illustrated in the upper pane in Fig. 1 right, by the variation in the intensity of the red shading). This fuzziness is mathematically represented by a probability of the possible values *m* can take, given the actual past *n*, i.e. *p*(*m*|*n*). The representation accuracy is quantified by the mutual information between the past *n* and the representation *m*, denoted by *I*(*n*; *m*), where the minimal value is 0 and the maximal value is the entropy *H*(*n*) (in the case of *m* = *n*). Another difference to note between the models is that although both models incorporate a compressed representation of the preceding sequence, the NOC model is simply defined by the minimal sufficient statistics, whereas the IB model is defined by the *-* log of probabilities.

### Fitting the EEG data

We assumed that the P300 response was linearly dependent on the IB subjective surprise (see the previous section for the definition of the IB subjective surprise). We extracted the P300 AUC feature on each trial as described in the electrophysiological analyses section. Independently, we calculated the subjective surprise for each trial for different *N* and *β* values; i.e., *S*^*β,N*^ (*z*_1_, *…z*_*n*_), where each such signal is called an IB surprise predictor. In order to avoid numerical problems caused by insignificant differences in surprise values, the surprise values were binned to bins of size 10^−4^ (using Matlab’s histc function). To characterize the best model we then calculated a linear regression fit between the single-trial AUC and the single-trial surprise values. Since there was an unbalanced distribution over the surprise values (by definition, higher surprise values are rarer), we used a weighted linear regression with inverse-probability weighting [65]. The inverse-probability 1*/p*(*s*) was calculated using the true asymptotic probabilities given by *p*(*y*_*t*+1_, *x*_*t*_) by summing over all probabilities with the same surprise value (rather than by the noisy empirical probabilities of the surprise values). In order to avoid multiple comparisons [25], P-values were calculated using a permutation test by randomizing the oddball sequence order within each block and then recalculating the surprise model of the permuted sequence. On 1000 permutations per subject, the maximal weighted-*R*^2^ from each permutation was extracted and used for the P-value calculation.

The optimal *N* in the NOC model was determined in a similar manner. For each *N,* an NOC predictor *n*_*t*_ was determined by calculating, for each tone in the sequence, the number of preceding tones of the opposite type in the preceding *N* tones (see the main text for more details). Then a weighted-*R*^2^ was calculated, where the weights were defined as for the case for the IB predictors, as inverse-probability weights using the true asymptotic probabilities. The probability *p*(*n*) in the predictors is given by *p*(*y*_*t*+1_ = 0, *x*_*t*_ = *n*) + *p*(*y*_*t*+1_ = 1, *x*_*t*_ = *N -n*).

### Significance testing

For the weighted linear regression in the single-trial, single-subject analysis, p-values were calculated using a 1000-fold permutation test by randomizing the sequence order within each block and then recalculating the IB or NO predictor of the permuted sequence. From each permutation the maximal weighted-*R*^2^ across all parameter space was extracted and used for the permutation test. For other linear regression analyses the reported p-values and F-statistics were calculated using Matlab’s built-in function *fitlm*.

## Acknowledgments

We thank Prof. Israel Nelken, Prof. Leon Deouell, Dr. Yoav Kessler and Dr. Tomer Fekete for many valuable discussions, Oren Alkoby, Mai Goffen, Erez Wolfson and Dr. Amjad Abu-Rmileh for assistance in preparing and running the experiments. N.T. and H.L.A. thank the Gatsby charitable foundation, and the Intel ICRI-CI for funding for the research.

## Supporting information

**S1 Fig.**
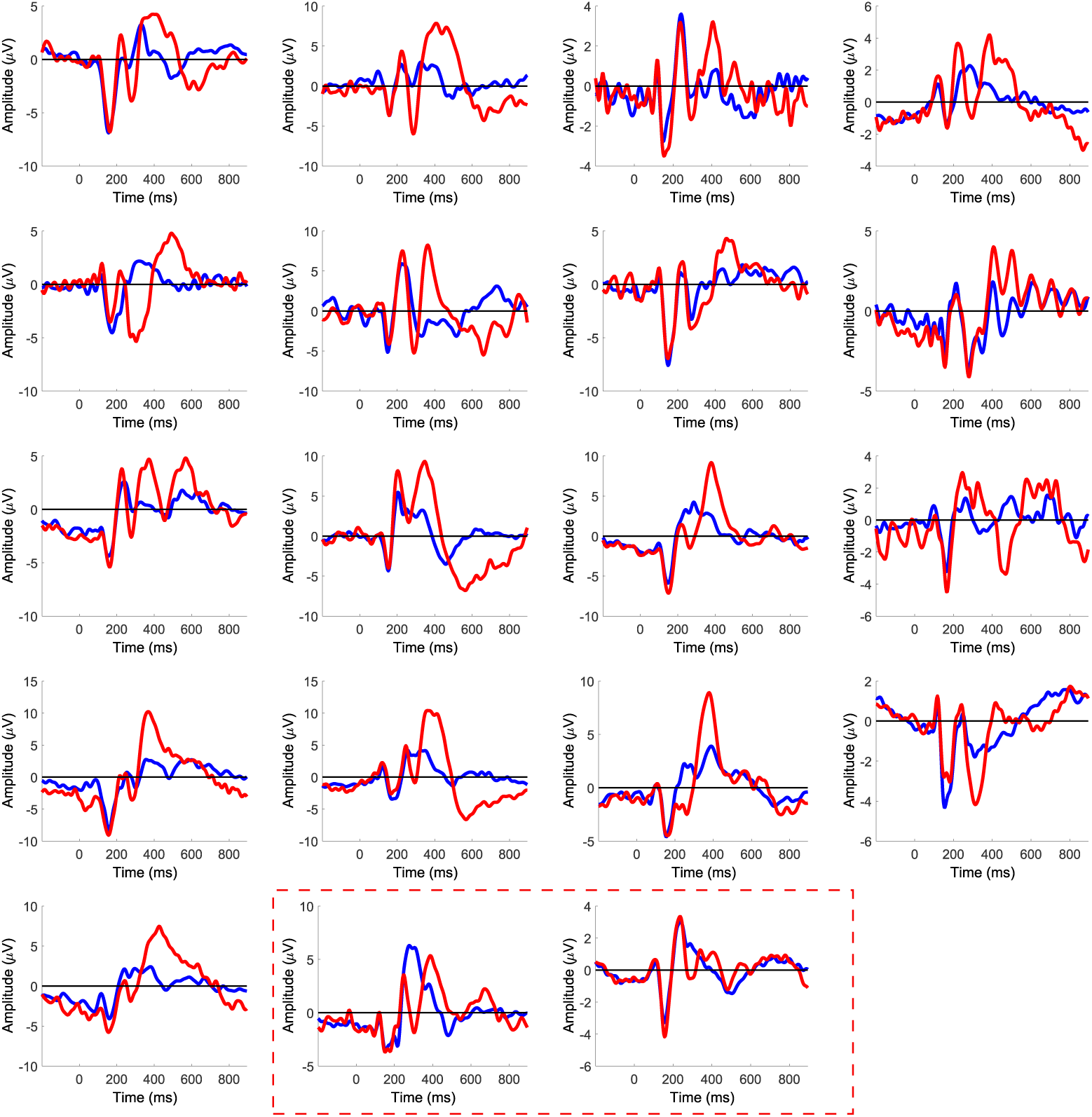
Single subject P300 ERPs. The P300 ERP of all subjects at electrode Cz is shown, each subject in a different plot. In red: the average of all oddball trials. In blue: the average of all standard trials. The last two plots marked by a dashed red rectangle correspond to two subjects where there was no clear difference between the oddball and standard curves and who therefore were omitted from the remainder of the analyses.

**S2 Fig.**
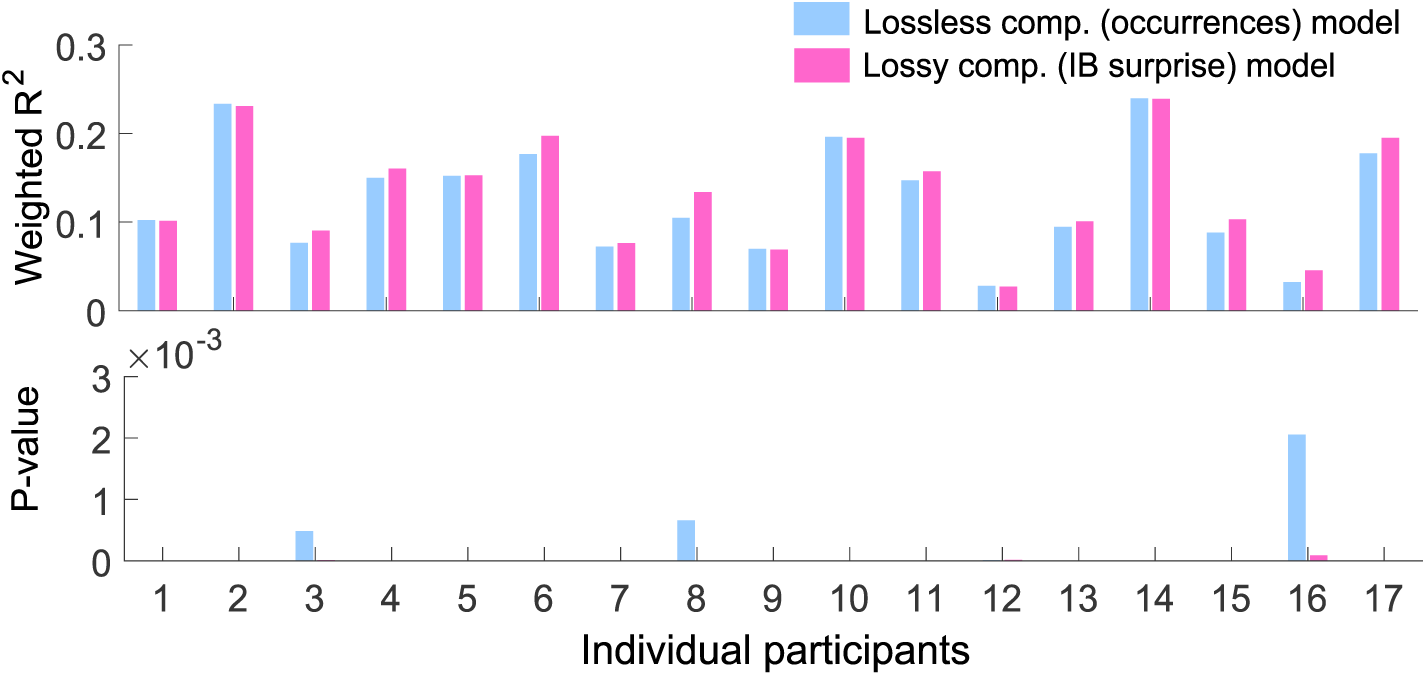
Subject-by-subject comparison of models performance. (**Top**) The weighted-*R*^2^ of the two optimal models is compared for each subject. Each pair of blue (NOC model) and magenta (IB model) bars depict a different subject. (**Bottom**) The p-value for each model is shown for each participant. The p-value was calculated with a 1000-fold permutation test. The significance of p-value=0 here is p¡0.001.

**S3 Fig.**
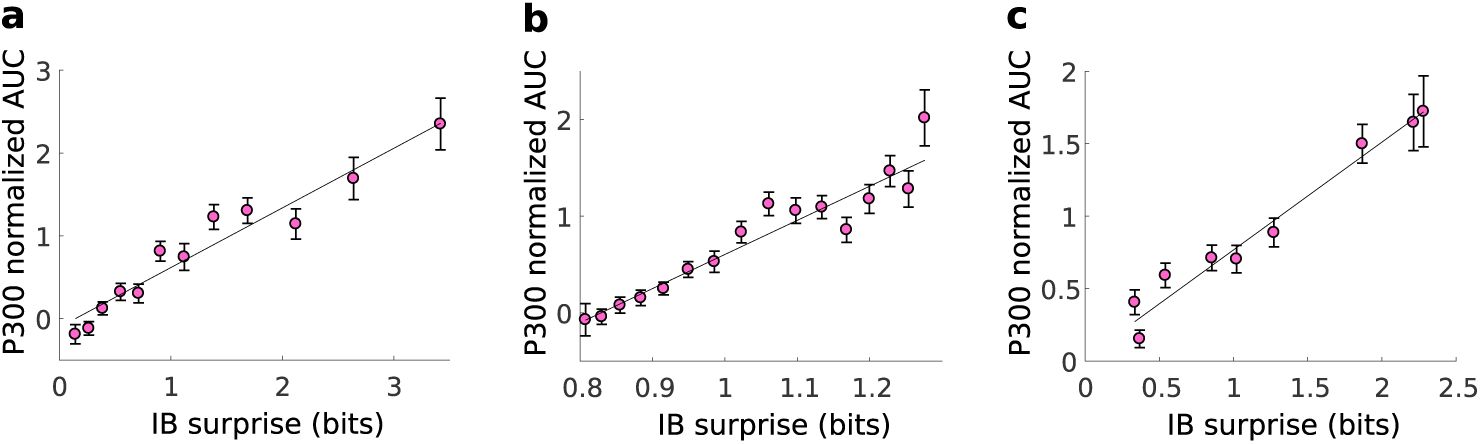
Single subject mean responses in the approximately sufficient compression model. Average normalized P300 AUC responses as a function of the IB surprise for three different subjects. The fitted parameters for each subject were: (a) *N* = 11, *β* = 48.33 (b) *N* = 15, *β* = 2.64 (c) *N* = 8, *β* = 11.29. The weighted-*R*^2^ values for the single-trials fit were: (a) 0.231 (b) 0.244 (c) 0.163. The *R*^2^ values for the fit of the mean values (the plotted line) were: (a) 0.939 (b) 0.907 (c) 0.966. The error bars indicate the SEM.

**S4 Fig.**
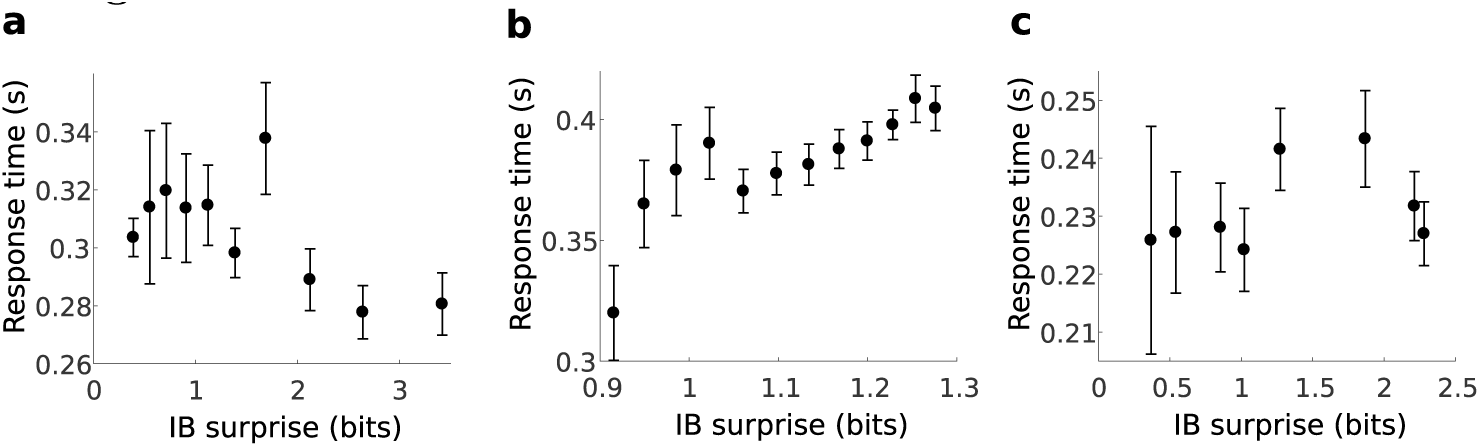
Single subject response time as a function of the IB surprise. Results for three subjects (same subjects as in supplementary S3 Fig), showing a non-linear and non-monotonous dependency of the RT on the IB surprise. This behavior was qualitatively different across subjects, showing that a multi-subject analysis is not straightforward for the RT and calls for a more complex model, presumably due to additional parameters affecting the response time. The surprise model parameters are as indicated in supplementary S3 Fig The error bars indicate the SEM.

**S5 Fig.**
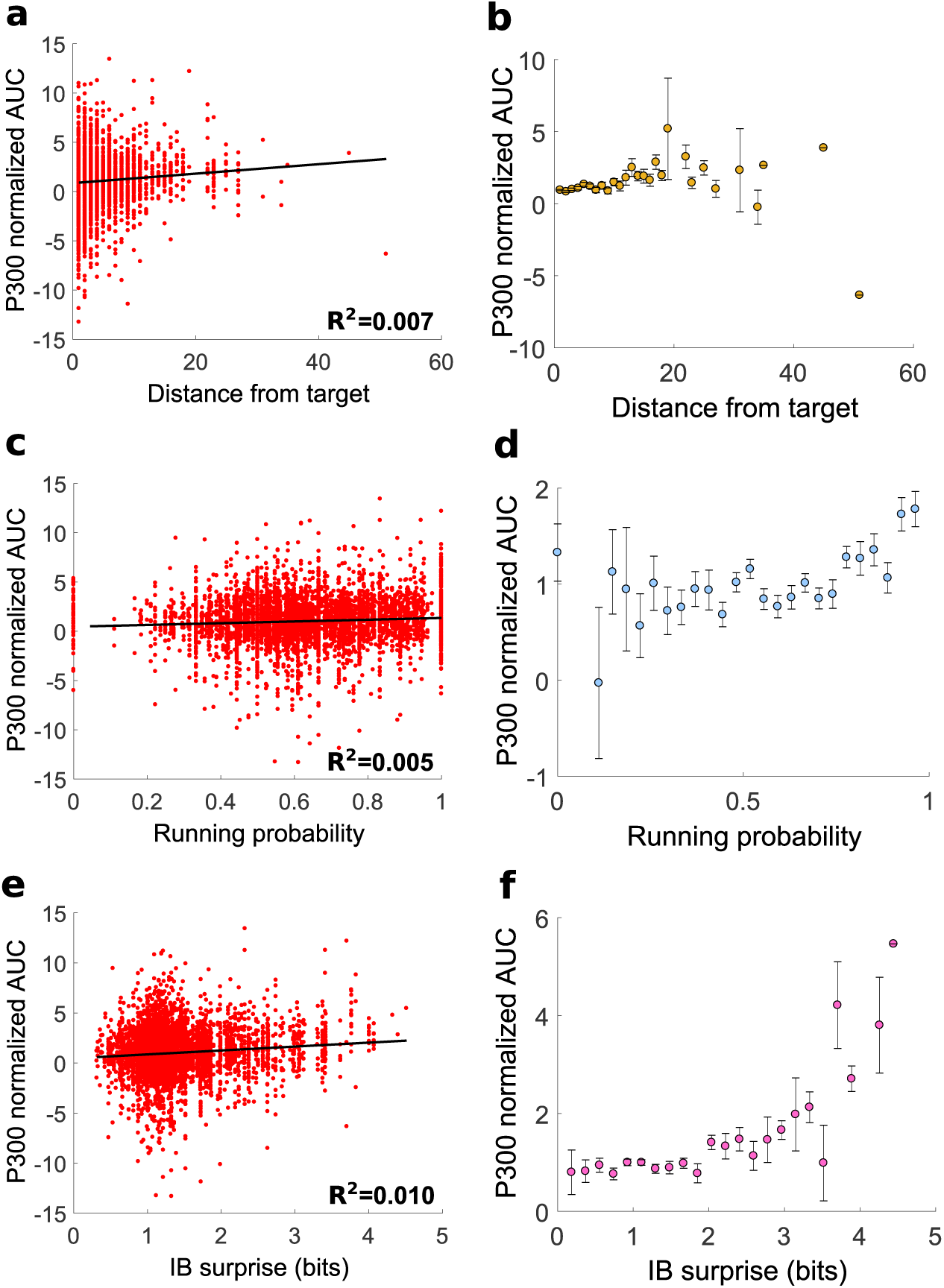
A comparison of the NOC and IB models with the distance-from-target model on oddball trials. (a) Single-trial and (b) average normalized P300 AUC responses to oddball tones as a function of the distance (in number of elements) from the last oddball in the sequence, for all subjects. *R*^2^ = 0.007, 5448 data points, error DOF = 5446, F-statistic vs. constant model: 37.3, p-value=1.06×10^−09^ (c) Single-trial and (d) average normalized P300 AUC responses to oddball tones as a function of the running probability, for all subjects. For each subject the best *N* was fitted as described in the main text. *R*^2^ = 0.005, 5492 data points, error DOF = 5490, F-statistic vs. constant model: 29.3, p-value=6.39×10^−08^ (e) Single-trial and (f) average normalized P300 AUC responses to oddball tones as a function of the IB surprise, for all subjects. For each subject the best *N* and *β* were fitted as described in the main text. *R*^2^ = 0.01, 5448 data points, error DOF = 5446, F-statistic vs. constant model: 55.3, p-value=1.21×10^−013^ The running probability and IB surprise were binned such that they have an identical number of values (28) on the x-axis as in (a). In all figures the error bars indicate SEM. Notice how the IB surprise shows a consistent increase in the AUC while explaining a large range of AUC responses.

### S1 Text Alternative surprise models for the P300

In the context of the oddball paradigm, Tueting, Sutton and Zubin [1] showed in 1970 that the P300 amplitude is affected by the oddball probability of the sequence (among other factors [2]). This was followed by other studies described below which showed dependence of the P300 on the preceding sequence of tones, in addition to the effect of the a-priori oddball probability. In 1976 an innovative study by Squires et al. [3] suggested a model of trial-by-trial expectancy to account for fluctuations in the P300 amplitude due to an auditory oddball sequence. This was an impressive study, but the model had several components and only considered the influence of up to five preceding elements. A study a year later by Duncan-Johnson and Donchin [4] compared the effect of the a-priori probability relative to the effect of the preceding tone and found that both factors contributed to the P300 amplitude independently. However, this only characterizes very short term memory effects (one preceding tone).

More recently, a model by Mars et al. [5] considered infinitely long sequences in the past. The surprise of each event is modeled as the minus log of the probability associated with each event given all preceding trials. The probability is estimated using a maximum likelihood estimate and assuming a prior with equally likely events (formally assuming a uniform Dirichlet prior over the oddball probabilities). This was the first work, to the best of our knowledge, to give a formal account of the surprise in single trials and associate it with single trial amplitudes of the P300. What is missing from this work, in our view, is the eventuality of inter-subject differences in the surprise model (all subjects have the same model with infinite memory length).

Finally, Kolossa et al. [6] suggested a predictive surprise model based on digital filtering, combining both Mars’ and Squires’ models and redefining them as three additive digital filtering processes. The surprise is modeled as the minus log of the probability of the next element, where the probability is given as a sum of three components: a short-term memory contribution depending on the number of oddball occurrences in the entire sequence with a strong decaying memory factor, a long-term memory contribution with a slower decay factor, and an alternation term which depends on a few preceding elements. Kolossa’s model parameters can be easily interpreted and connected to memory parameters; however, as Squires’ model it seems to have a relatively large number of components and parameters.

Kolossa et al. thoroughly compared the above models [6] and also drew the attention to the difference between models of predictive surprise and models of Bayesian surprise. In models of predictive surprise the probability for the next element is estimated in each trial and the surprise in each trial is modeled as the minus log of this probability. Models of Bayesian surprise model the surprise as the revision in the internal probability distribution over the possible elements after each trial. This is the distance between the two estimated distributions, before and after observing each element. This can be quantified, for example, using the *D*_*KL*_ distance between the distributions. An example of a Bayesian surprise model was given by Ostwald et al. [7] for the somatosensory system under an oddball paradigm. However, Mars et al. [5] tested a Bayesian surprise model on their P300 data as an alternative model and found their predictive surprise model to give better results.

As we show in the main text, the dependency on the number of oddball occurrences is observed for a good theoretical reason: given a memory length, in the oddball paradigm the number of oddball occurrences in the preceding sequence is a minimal sufficient statistic [8] to predict the next tone. The models mentioned above are all dependent on this number, but Squires’ and Kolossa’s models contain more information about the exact sequence which is both unnecessary theoretically for efficient processing of the oddball sequence, and also do not seem to have a significant advantage in explaining the P300 data, as shown in Kolossa et al. Mars’ model, on the other hand, may lose important information by unifying all subjects in a single model.

It is also worth noting a related predictor for the P300 amplitude known as the target-to-target interval [9]; i.e., the number of non-target elements preceding the target element. This predictor considers only target trials and was used to analyze mean responses. A comparison with this model is shown in supplementary S5 Fig.

